# Genomic characterization of a pathogenic isolate of *Saccharomyces cerevisiae* reveals an extensive and dynamic landscape of structural variation

**DOI:** 10.1101/2021.08.20.457152

**Authors:** Lydia R. Heasley, Juan Lucas Argueso

## Abstract

The budding yeast *Saccharomyces cerevisiae* has been extensively characterized for many decades and is a critical resource for the study of numerous facets of eukaryotic biology. Recently, the analysis of whole genome sequencing data from over 1000 natural isolates of *S. cerevisiae* has provided critical insights into the evolutionary landscape of this species by revealing a population structure comprised of numerous genomically diverse lineages. These survey-level analyses have been largely devoid of structural genomic information, mainly because short read sequencing is not suitable for detailed characterization of genomic architecture. Consequently, we still lack a complete perspective of the genomic variation the exists within the species. Single molecule long read sequencing technologies, such as Oxford Nanopore and PacBio, provide sequencing-based approaches with which to rigorously define the structure of a genome, and have empowered yeast geneticists to explore this poorly described realm of eukaryotic genomics. Here, we present the comprehensive genomic structural analysis of a pathogenic isolate of *S. cerevisiae*, YJM311. We used long read sequence analysis to construct a haplotype-phased, telomere-to-telomere length assembly of the YJM311 diploid genome and characterized the structural variations (SVs) therein. We discovered that the genome of YJM311 contains significant intragenomic structural variation, some of which imparts notable consequences to the genomic stability and developmental biology of the strain. Collectively, we outline a new methodology for creating accurate haplotype-phased genome assemblies and highlight how such genomic analyses can define the structural architectures of *S. cerevisiae* isolates. It is our hope that through continued structural characterization of *S. cerevisiae* genomes, such as we have reported here for YJM311, we will comprehensively advance our understanding of eukaryotic genome structure-function relationships, structural diversity, and evolution.

## Introduction

*Saccharomyces. cerevisiae* has emerged as a robust system for the study of population genetics and species evolution^1,2^, due in large part to concerted efforts of yeast geneticists to collect, curate, and sequence >1000 natural isolates^3,4^. This wealth of whole genome sequencing (WGS) data, most of which has been generated using short read technologies, has provided insights critical to numerous aspects of *S. cerevisiae* biology including genome evolution, population diversity, and genotype-phenotype relationships^3,5-7^. Collectively, these survey-level genome investigations have demonstrated that this model organism has a complex and fascinating evolutionary history^8,9^, and have highlighted the tremendous genomic diversity that can exist within a single species^4,10,11^. Whereas short read WGS data is ideal for assessing genomic heterozygosity and allelic variation between individuals, long read sequencing approaches can reveal the structural architecture of the genome^12^. The advent of single molecule long read WGS technologies (*e*.*g*., PacBio, Oxford Nanopore Minion) has enabled researchers to begin exploring the structural contributions to the overall genomic variation existing within the *S. cerevisiae* species. The roles of structural genomic architecture and variation in the function and evolution of the *S. cerevisiae* genome are likely to be significant, and continued investigation into these genomic features are needed to advance our understanding of the life history and diversification of the species.

Here we report an in-depth structural genomic analysis of the strain YJM311. YJM311 is a clinically derived isolate that displays several pathogenically relevant phenotypes^13-15^. One such phenotype, called the colony morphology response, enables YJM311 cells to transition from a unicellular growth state to a cooperative ‘multicellular’ growth state in response to environmental changes in carbon source (*i*.*e*., glucose) availability^16,17^. On media rich in glucose, YJM311 cells form a typical smooth and simple colony architecture, but when plated on glucose-poor media, cells form complex colonies with a highly patterned organization^16-18^. Previously, studies using YJM311 as a model with which to investigate the genetic regulation of the colony morphology response demonstrated that allelic variations at numerous sites in the diploid genome modulate this complex phenotypic trait^17^. Together, Granek *et al*., found that most genes involved in this cellular behavior encode protein products whose activities regulate highly conserved environmental stress pathways^17^. Recently, we observed that some YJM311 cells stochastically switched between these simple and complex phenotypic states, even when grown in identical conditions. While investigating the sources of this phenotypic variation^18^, we also took the opportunity to explore the structural composition the YJM311 genome. By combining the of strengths short- and long read WGS analyses, we assembled the fully phased heterozygous diploid genome of YJM311 and explored the myriad structural complexities comprising the genomic architecture of this strain.

## Results

### Structural features of the YJM311-3 genome

Before initiating our studies of YJM311^18^, we first assessed the homogeneity of our stock by purifying individual clones for comparative karyotype analysis using pulsed-field gel electrophoresis (PFGE)(*i*.*e*., YJM311-1 through YJM311-7)(Fig. 1). From this, we discovered that our YJM311 stock consisted of a mixture of karyotypically similar, but discernible subpopulations. Comparison of these clones indicated that YJM311-1/YJM311-2, YJM311-3/YJM311-7, YJM311-4/YJM311-6, and YJM311-5 represented four distinct karyotypes. Because the molecular karyotypes of the YJM311 subclones differed substantially from that of the reference genome strain S288c, it was difficult to distinguish which chromosomes in YJM311 carried the SVs visible by PFGE analysis (Fig. 1). To determine the chromosomal identities of the individual bands in the YJM311 karyotype, we used long read WGS analysis to assemble the fully phased diploid genome of YJM311-3. We chose to assemble the genome of this specific subclone because individual chromosomal bands were better resolved in the molecular karyotype of YJM311-3/YJM311-7 than in the karyotypes of the other subclones (Fig. 1). We also used this subclone to characterize the sources of stochastic phenotype switching displayed by YJM311^18^.

**Figure 1.**
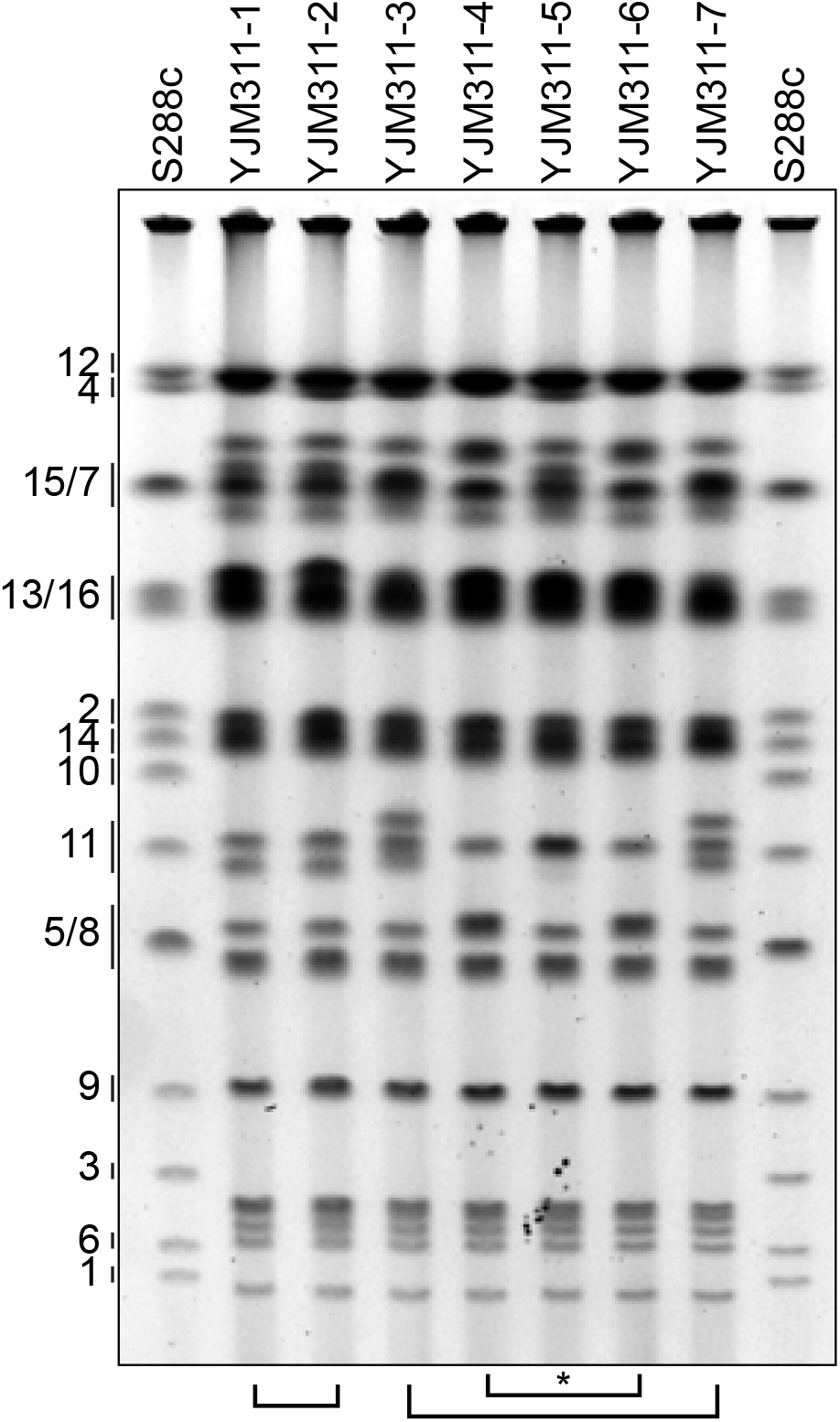
The YJM311 stock consists of karyotypically distinct subpopulations. Shown is a PFGE molecular karyotype of the reference genome strain S288c and a sampling of seven independent clones recovered from the stock population of YJM311. Numbers along the left delineate the chromosome identities represented by the bands present in the S288c molecular karyotype. Brackets at the bottom of the gel denote YJM311 clones that share the same karyotype. Asterisk denotes that YJM311-5 has a unique molecular karyotype.

To assemble the phased diploid genome, we generated Oxford Nanopore Minion long reads, phased these reads by haplotype using the long read variant caller Longshot^19^, and performed separate *de novo* genome assemblies using each set of haplotype-phased long reads (See Materials and Methods)(Fig. S1). Collectively, this assembly method produced 32 chromosome-length contigs, each of which corresponded to one of the homologous chromosomes comprising the YJM311 diploid genome. To our knowledge, the only mis-assembled genomic region in this assembly is the long and highly repetitive ribosomal DNA (rDNA) array, located on Chr12. The contigs corresponding to the two homologs of Chr12 contain rDNA arrays 75kb and 85kb in length, respectively. This is almost certainly incorrect, as studies have determined that the length of the rDNA array in *S. cerevisiae* can reach ∼1,350kb in length^20^. Indeed, manual analysis of reads spanning the rDNA array identified long reads containing 12 rDNA repeats. These reads were among the longest in our read set (∼130kb) yet still failed to span the complete length of the rDNA array. To normalize the structure of the rDNA array similar to the approach used to represent the rDNA array in the S288c reference, we arbitrarily edited the Chr12 contigs such that they each carry 2 rDNA repeat units.

Analysis of the YJM311-3 genome assembly revealed pervasive structural variation which differed from the S288c reference genome and between the homologous pairs of YJM311 chromosomes. Much of this interhomolog structural variation constituted heterozygous Ty element insertions and sub-telomeric variations. Collectively, the *de novo* assembly of the YJM311-3 genome contains 65 Ty elements, most of which differ between homologs in terms of genomic position, orientation, and Ty family (Fig. 2A, yellow arrows). Additionally, many chromosome pairs differ in structure at the sub-telomeres. Several chromosomes, such as Chr5*a*, Chr7*b*, and Chr11*a*, contain sizeable tandem arrays of sub-telomeric Y’ element sequences (Fig. 2B, turquoise arrows). In the case of the left telomere of Chr5*b*, we were unable to deduce the complete terminal structure, as our read set lacked reads long enough to span the whole Y’ array and reach the telomeric consensus repeats. Thus, we can only infer that the left sub-telomere of Chr5*b* has a Y’ array containing at least 9 tandem Y’ elements. In the case of the right telomere of Chr7b, we identified reads long enough to establish the presence of a Y’ array consisting of 12 consecutive tandem repeat units followed by consensus telomeric repeats.

**Figure 2.**
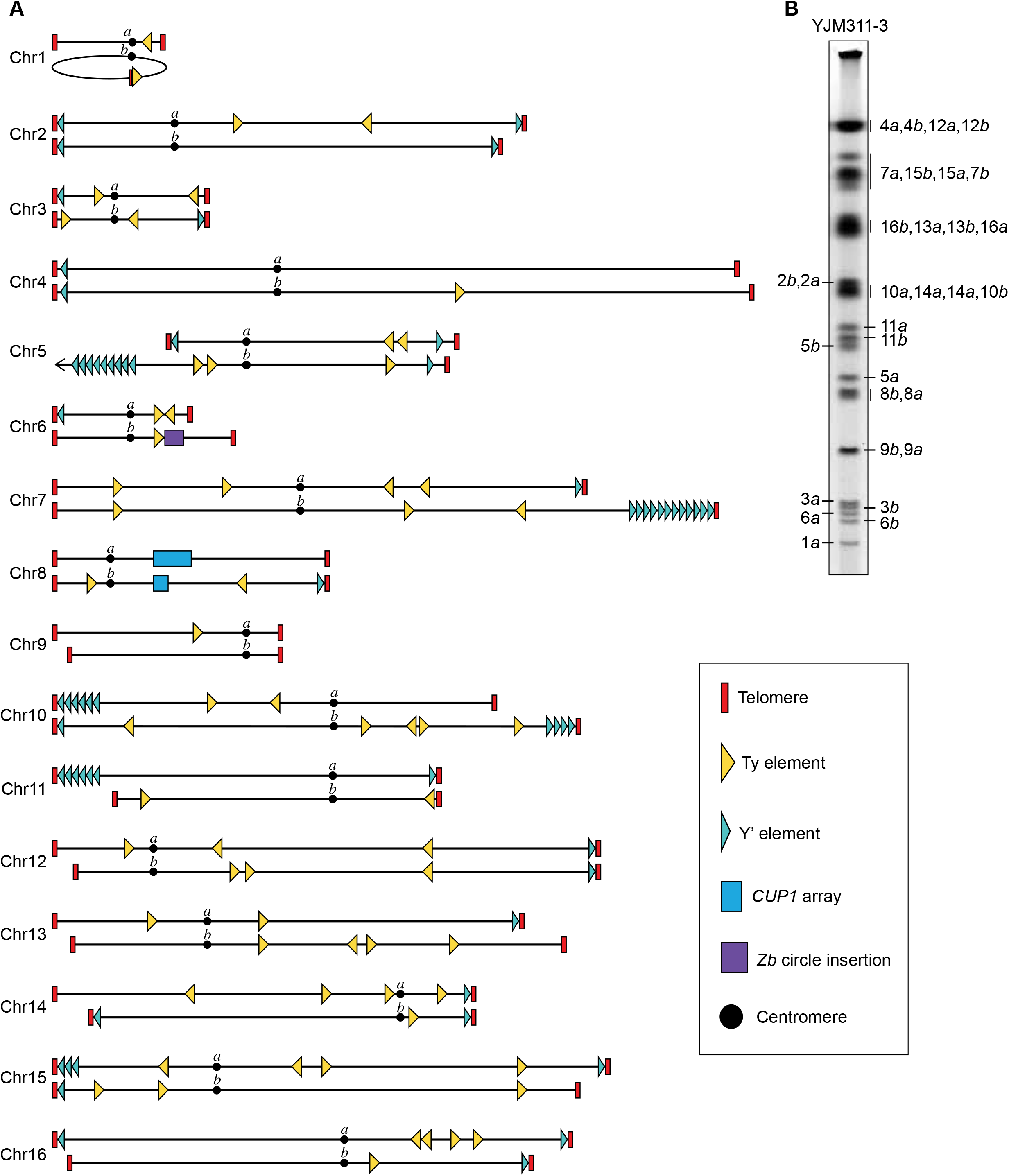
The structural architecture of the haplotype-phased diploid genome of YJM311-3. **A**. A schematic of the *de novo* assembly of the YJM311-3 genome. Structural features discussed in the text are denoted in the boxed legend at right. A complementary map of the genomic distribution of hetSNPs is shown in Figure S1. **B**. The PFGE molecular karyotype of YJM311-3/YJM311-7 from Fig. 1 annotated with the chromosomal identities of specific bands.

Previous studies have demonstrated that the architecture of the *CUP1* locus can vary greatly between strains^21^. Indeed, when we inspected the different structures of the *CUP1* arrays present on Chr8, we found that in YJM311-3, each homolog of Chr8 carries arrays which differ in number and type of repeat unit (Fig. 3A). Chr8*b* has a *CUP1* array ∼8kb in length and composed of seven Type 3 repeats. The Type 3 repeat unit consists of the *CUP1* gene, the *RUF5* noncoding RNA, and the replication origin *ARS810*^21^. Chr8*a* has a substantially longer array (∼24kb) consisting of 14 repeat units. The three most centromere-proximal repeats are Type 3 units identical to those found on Chr8*b*, and the remaining 11 repeats are Type 1 units^21^. Type 1 repeat units, which comprise the *CUP1* array of the S288c reference genome, consist of *CUP1, RUF5, ARS810*, and the open reading frame *YHR054C*. The presence of both Type 3 and Type 1 repeat units in the Chr8*a* hybrid array demonstrates that while the ancestral homologs of YJM311 likely carried arrays composed of distinct repeats, interhomolog recombination has contributed to the further diversification of the *CUP1* locus in this strain.

**Figure 3.**
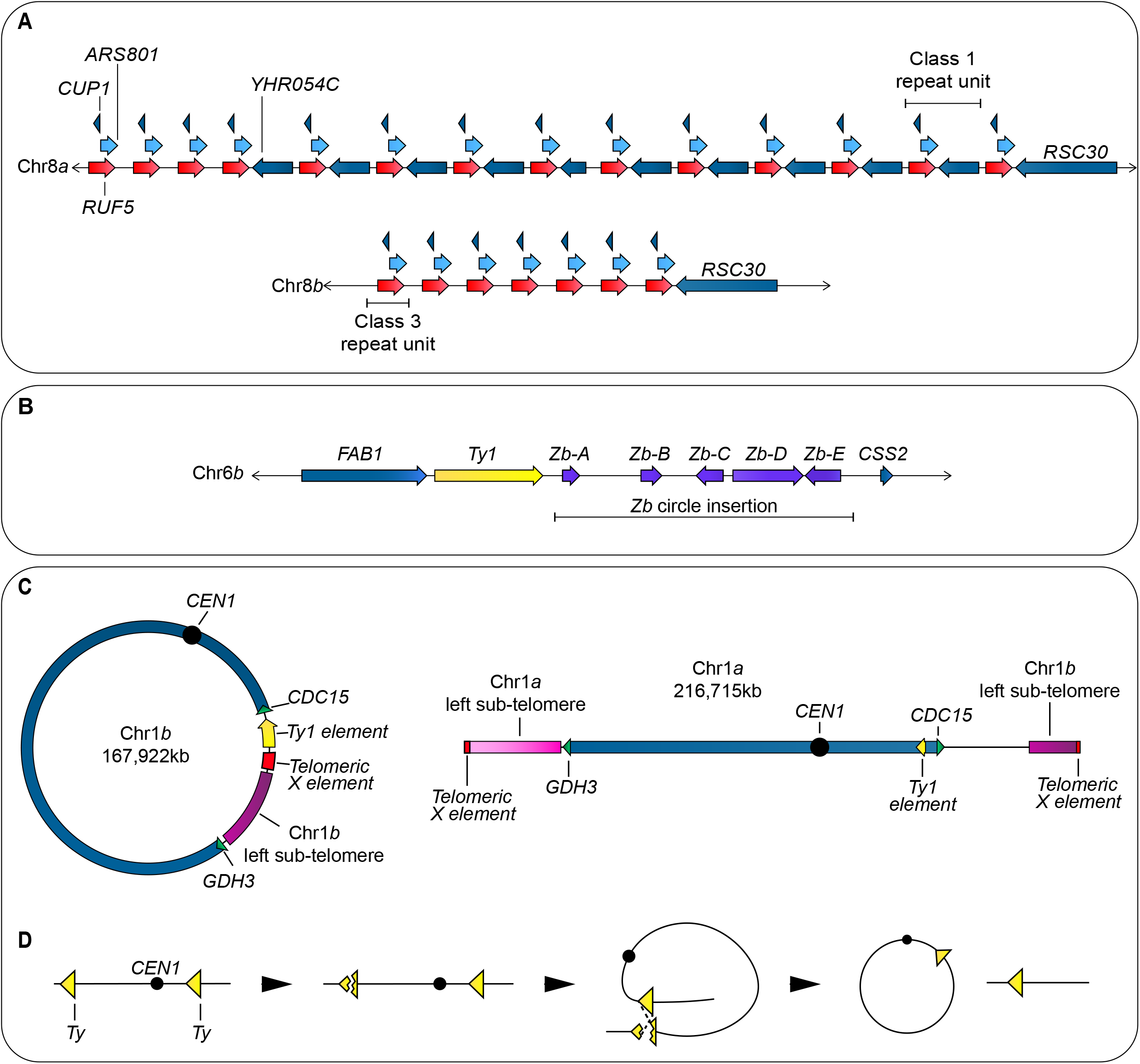
Detailed analyses of structurally complex regions in the YJM311-3 genome. **A**. A schematic detailing the organization of the *CUP1* arrays located on the two homologs of Chr8. **B**. A schematic detailing the hemizygous insertion of the *Zygosaccharomyces bailii* circle on the right arm of Chr6*b*. **C**. Schematics detailing the structures of the circular (Chr1*b*) and linear (Chr1*a*) homologs of Chr1. **D**. A simple model explaining the formation of a circular chromosome through mitotic recombination between Ty1 elements located on opposite sides of the centromere.

We discovered a ∼17kb region on the right arm of Chr6*b* that failed to align to the *S. cerevisiae* reference genome, and identified this sequence, which is distal to a Ty1 element adjacent to the *FAB1* gene, as a set of 5 genes (*ZbA-ZbE*) derived from an extrachromosomal circle harbored by *Zygosaccharomyces bailii* (Fig. 3B)^22,23^. Acquisition of this extrachromosomal sequence by *S. cerevisiae* is posited to have occurred through a horizontal gene transfer event, and numerous distinct insertions of this sequence have been identified in the genomes of other strains, primarily in those isolated from wine-making environments^23^. The specific insertion of the *Z. bailii* circle at this locus on Chr6 appears to be shared by other wild isolates of *S. cerevisiae*, as we recently identified an identical insertion in one of the Chr6 homologs in the diploid genome of the industrial strain JAY270^24^.

Perhaps the most striking SV present in the YJM311-3 genome is the circularization of one homolog of Chr1 (Fig. 3C, Chr1*b*). Despite its smaller size, the circular nature of Chr1*b* precludes it from being resolved using PFGE approaches and for this reason we were only able to visualize the linear homolog of Chr1 (Chr1*a*) in the molecular karyotype analysis of all YJM311 subclones (Fig. 1A, Fig. 2B). Chr1*a* has also undergone a rearrangement which renders it an isochromosome, with sub-telomeric sequence derived from the left telomere on each end of the chromosome (Fig. 3C). The structure and hetSNPs present at each sub-telomeric region of Chr1*a* differ from one another, indicating that Chr1*a* did not become an isochromosome through recombination with itself, but through a recombination event with Chr1*b* or the linear precursor of Chr1*b*. The circular Chr1*b* molecule contains a telomeric X element directly adjacent to a Ty1 element distal to the essential gene *CDC15*. While numerous possibilities exist, one of the simplest models explaining the circularization of Chr1*b* would be recombination between the Ty1 element adjacent to *CDC15* and another Ty1 element present near the left telomere of Chr1*b* (Fig. 3D). Recombination between these Ty1 elements would produce a circular molecule carrying the centromere, and an acentric molecule, which would be lost over time.

### Y’ mediated telomeric variations underlie the karyotypic differences between YJM311 subclones

To determine the basis of the structural variations (SVs) existing between the karyotypically distinct subclones of YJM311 (Fig.1), we performed short read WGS analysis of YJM311-1, YJM311-3, YJM311-4, YJM311-5, and YJM311-6. With the exception of several large tracts of homozygosity existing on Chr2, Chr5, Chr12, and Chr15, the YJM311genome was previously found to be highly heterozygous (∼40,000 hetSNPs)(Fig. S2A)^8^. We determined the distribution of hetSNPs in the genomes of each YJM311 subclone, with the expectation of identifying unique *de novo* tracts of homozygosity arising from mitotic recombination which would explain the chromosome size polymorphisms reflected in our PFGE analysis (Fig. 1). Surprisingly, we found that the hetSNP profiles of each YJM311 derivative were identical to one another and to that of earlier published analysis (Fig. S2B)^8^. From this, we concluded that the SVs visible by PFGE analysis likely reflected genomic alterations not easily captured by short read WGS, such as reciprocal translocations or SVs present in repetitive regions of the genome (*e*.*g*., Ty element insertions, telomeres, and sub-telomeric regions).

Using the phased genome assembly of YJM311-3, we were able to deduce most of the chromosomal identities of the bands in the PFGE karyotypes of the YJM311 subclones. From this, it became clear that the most variable bands in the karyotypes of the YJM311 subclones corresponded to Chr5*b* and Chr11*a* (Fig. 1, Fig. 4A). We generated Minion long reads from YJM311-1, YJM311-4, and YJM311-5 and aligned the reads from each subclone to the phased YJM311-3 genome assembly so as to determine the structural basis of the band shifts reflected in the PFGE gel, with a particular focus on Chr5*b* and Chr11*a*. Interestingly, we did not detect any reciprocal crossovers relative to the YJM311-3 genome and instead found that the sub-telomeric regions, specifically the Y’ arrays, were the only parts of Chr5*b* and Chr11*a* which differed between the YJM311 subclones. Our analysis of these distinct Y’ arrays was limited by to the extensive length of Y’ arrays on Chr5*b* (>58kb) and Chr11*a* (∼40kb), as it was difficult to identify reads unique to each chromosome that spanned the full Y’ array and terminated with telomere consensus repeats. However, using our molecular karyotype analysis and the longest reads generated for each subclone, we were able to infer the different sub-telomeric structures in YJM311-1, YJM311-4, and YJM311-5 relative to YJM311-3.

**Figure 4.**
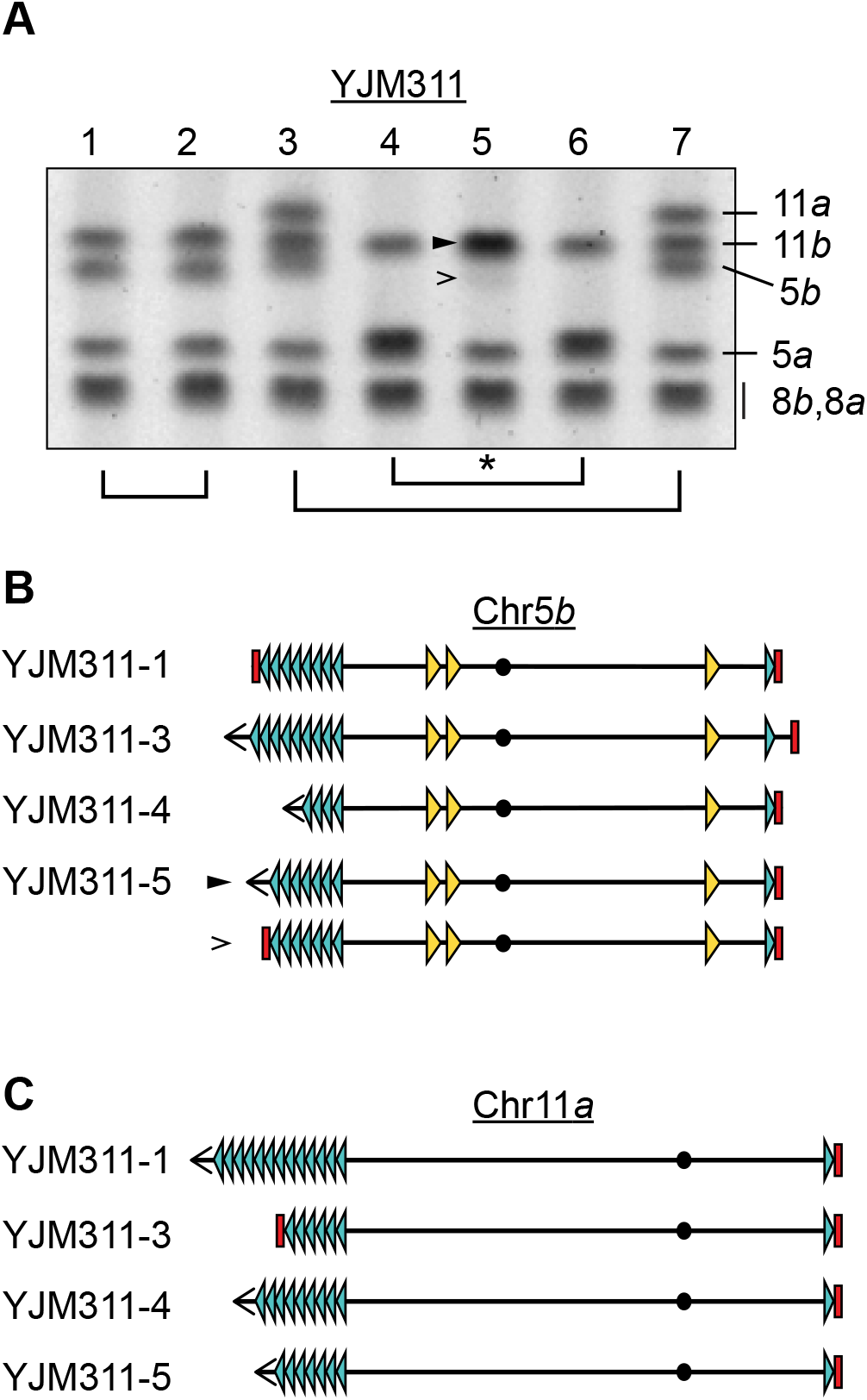
Dynamic rearrangements of sub-telomeric Y’ arrays underlie the karyotypic diversity of YJM311 subclones. **A**. A cropped image of the PFGE molecular karyotypes from Fig. 1 highlighting the size differences of Chr5*b* and Chr11*a* among YJM311 subclones. Solid and open arrowheads in the karyotype of YJM311-5 denote the two different species of Chr5*b*. The brackets and asterisk below are the same as in Fig. 1. **B**. Schematics of the structural characteristics of Chr5*b* in the denoted subclones. **C**. Schematics of the structural characteristics of Chr11*a* in the denoted subclones. Symbols in B and C are the same as in Fig. 2.

First, we analyzed the sub-telomeric structures of Chr5*b* (Fig. 4A, B). Our analysis demonstrated that relative to YJM311-3, YJM311-1 harbors a Chr5*b* containing fewer Y’ units at the left telomere and as well as a deletion of the sequence distal to the Y’ element at the right telomere. This deletion is likely the result of non-allelic recombination between the Y’ element on the right arm of Chr5*b* and a Y’ element on another chromosome. YJM311-4 harbors a significantly smaller Chr5*b* than YJM311-3, consistent with a major contraction of the left telomeric Y’ array combined with the same short right telomere as in YJM311-1. YJM311-5 appears to consist of a mosaic population of cells carrying one of two different Chr5*b* molecules. The majority of cells harbor the larger Chr5*b* species which co-migrates with Chr11*b*. This molecule likely harbors an expanded Y’ array at the left telomere. A minority of cells harbor a Chr5*b* which is smaller than that of YJM311-3 (faint band in Fig. 4A; “>“). Individual single molecule read analysis confirmed that this molecule has a shorter Y’ array on the left telomere.

We also analyzed the variable structures of Chr11*a* among YJM311 subclones (Fig. 4A, C). We found that relative to YJM311-3, the Chr11*a* molecules harbored by YJM311-1, YJM311-4, and YJM311-5 all contain a greatly expanded Y’ array at the left telomere. Indeed, YJM311-3 was the only subclone for which we detected reads that mapped to Chr11*a* and terminated with telomeric repeats. Read analysis for YJM311-1 indicated that the Y’ array at the left telomere consists of at least 13 Y’ elements. For this reason, the Chr11*a* molecules harbored by YJM311-1, YJM311-4, and YM311-5 co-migrate with the Chr2, Chr10, and Chr14 homologs in the PFGE (Fig. 1A). Collectively, this analysis revealed that the structures of sub-telomeric Y’ arrays explain the structural variation visible by PFGE between YJM311 subclones and further highlights the plasticity of these regions of the YJM311 genome.

### Chr1b is the source of poor spore viability in YJM311

The original characterization of YJM311 by McCusker and colleagues demonstrated that similar to other pathogenic isolates of *S. cerevisiae*, YJM311 sporulated with high efficiency yet produced spores with poor viability (36.1%)^14^. In that seminal report, the authors noted that there was no straightforward explanation for this poor viability, as tetrads did occasionally produce three or four viable spores^14^. Given that our phased genome assembly contained new insights into the structural architecture of the YJM311 genome, we used it to reinvestigate the source of spore inviability in YJM311. We hypothesized that the production of inviable spores could be attributed to problematic meiotic recombination between the linear and circular homologs of Chr1. In a simplistic model where only one chromatid from each homologous pair is involved in meiotic recombination, a single crossover event between a Chr1*a* chromatid and a Chr1*b* chromatid would result in the formation of a dicentric recombinant product (Fig. 5A)^25^. Because it contains two centromeres, this dicentric recombinant chromosome would be erroneously segregated or broken during meiosis I, and this would necessarily compromise the viability of the resulting spores. In the latter case, in which the dicentric chromosome is broken during anaphase I, two spores would go on to inherit intact non-recombinant copies of Chr1 during meiosis II, but the other two spores would inherit the broken fragments of the dicentric chromosome and would therefore be inviable (Fig. 5B). In contrast, a double crossover event between the same chromatids of Chr1*a* and Chr1*b* could result in the formation of recombinant linear and circular chromosomes and would not impact the viability of the spore products (Fig. 5B). During meiosis, small chromosomes such as Chr1 experience higher frequencies of meiotic crossovers per kilobase than larger chromosomes^26,27^, and crossover analysis in a hybrid diploid strain with two linear homologs of Chr1 demonstrated that 86% of spores inherited a recombinant Chr1 chromatid^28^. Therefore, in the likely scenario that Chr1 chromatids become involved in multiple meiotic crossovers, the formation of additional acentric and dicentric products would result in increased aneuploidization of Chr1 and further reductions in spore viability. Based on this model, we predicted that viable spores derived from YJM311-3 should be depleted for recombinant Chr1 products, specifically those with an odd number of crossovers.

**Figure 5.**
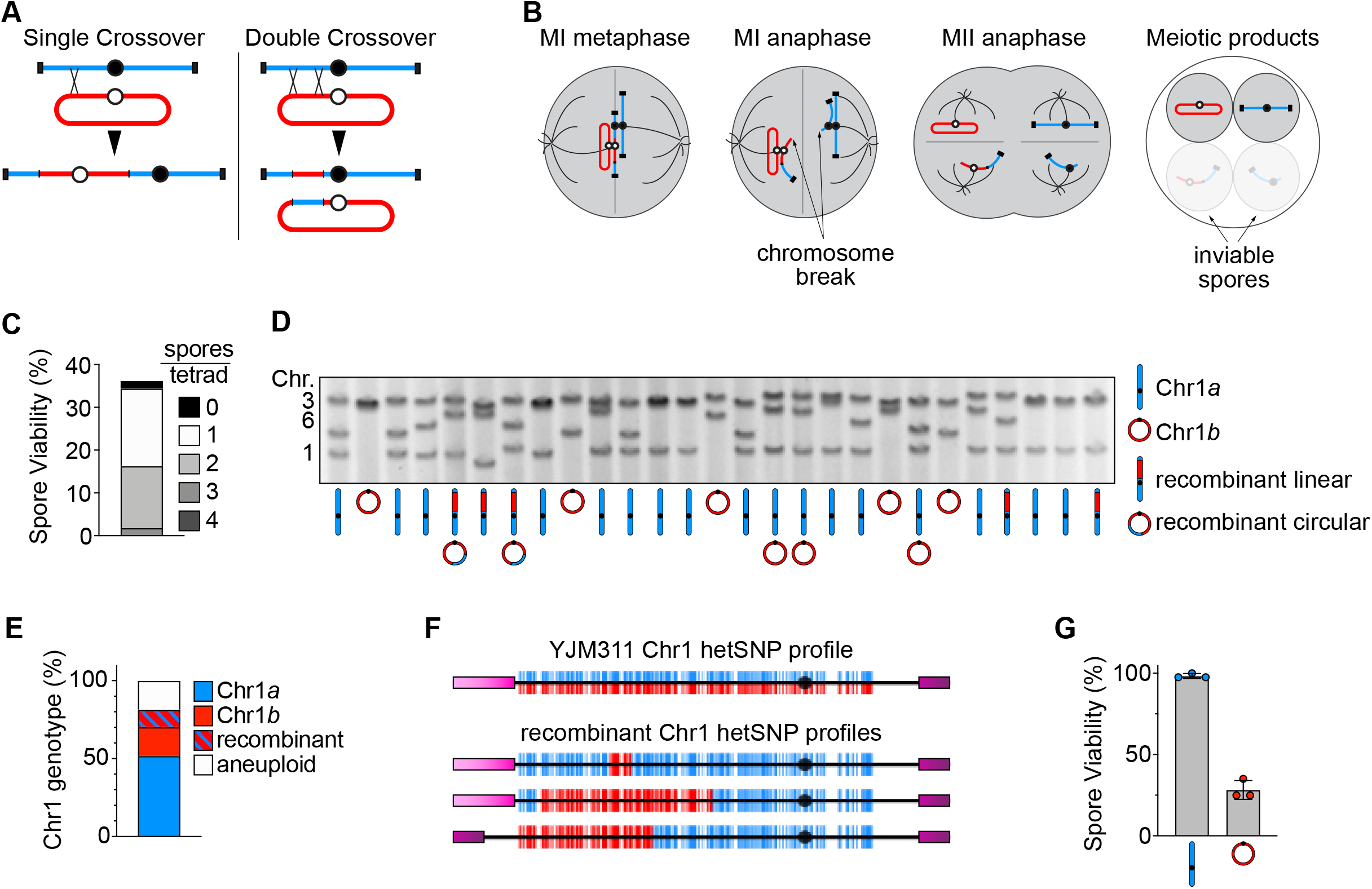
Meiotic errors arising from recombination between Chr1*a* and Chr1*b* are responsible for the poor viability of YJM311 spores. **A**. A schematic outlining the recombination products formed by single and double crossovers between circular and linear chromatids. The other two chromatids from each replicated homolog are not depicted. **B**. A model of meiotic chromosome segregation following the single crossover event between a circular and linear chromosome as shown in A. Segregation of a recombinant dicentric chromatid during MI anaphase would likely lead to formation of an anaphase bridge or broken chromosome. **C**. The percent total spore viability, and distribution of viable spores per tetrad in YJM311-3. **D**. A cropped image of the PFGE karyotype analysis of 27 viable spores derived from YJM311-3. This region of the karyotype shows bands corresponding to Chr1, Chr6, and Chr3. The genotypes of Chr1 as deduced by WGS analysis for each spore are shown as schematics below the cropped image. **E**. The percentage of viable spores analyzed in D. that harbor the denoted genotypes of Chr1. Blue chromosome, non-recombinant Chr1*a* (linear); Red chromosome, non-recombinant Chr1*b* (circular); Striped blue/red chromosomes, recombinant Chr1. **F**. HetSNP profiles of YJM311 Chr1 and recombinant Chr1 molecules harbored by three viable spores. In each profile, the black circle denotes the centromere, red lines represent the hetSNP identity of Chr1*a*, blue lines denote the hetSNP identity of Chr1*b*. Magenta and purple boxes denote the heterozygous but allelic telomeres of Chr1*a*. **G**. The viability of spores derived from the diploidized segregants from YJM311-3 harboring either Chr1*a* or Chr1*b*. For each genotype, 10 tetrads (40 spores) were dissected from each of three unrelated diploidized spores harboring either only Chr1*a* or Chr1*b*. Error bars depict the mean spore viability of the three separate dissection experiments.

We tested this prediction using Illumina WGS and PFGE analysis to characterize the structure of Chr1 in 27 viable spores recovered from YJM311-3 (Fig. 5D-F). We dissected 20 tetrads from YJM311-3 and calculated the percent spore viability and the number of viable spores per tetrad (Fig. 5C). Consistent with the earlier analysis performed by McCusker *et al*., we observed similarly poor spore viability (36.25%) with most tetrads producing only one or two viable spores. PFGE and hetSNP analysis of the viable spores revealed that the majority of these spores harbored non-recombinant Chr1 molecules (70.4%) and were particularly enriched for inheritance of a non-recombinant linear Chr1*a* chromatid (51.9%) (Fig. 5D, E). Several spores harbored aneuploidies of Chr1, an outcome consistent with the predicted complications arising from meiotic recombination between multiple Chr1 chromatids during meiosis I. Of the spores that did carry a recombinant copy of Chr1 (Fig. 5F), all were linear molecules that displayed a SNP profile consistent with a double crossover event (Fig. 5F).

YJM311 has functional alleles of the *HO* gene encoding the homing endonuclease and is therefore homothallic. Upon germination, homothallic haploid spores are competent to switch mating type and haplo-self to form homozygous diploids capable of sporulating again^29^. We used the diploidized spores from YJM311-3 to determine whether spores carrying two copies of Chr1*a* exhibited rescued spore viability relative to spores carrying two copies of Chr1*b*. Indeed, homozygous diploids harboring Chr1*a* displayed a complete restoration of spore viability to nearly 100% (Fig. 5G, n=120 spores derived from three independent diploids for each genotype). In contrast, homozygous diploids carrying Chr1*b* displayed poor spore viability similar to that exhibited by the parental YJM311-3 (Fig. 5G). This effect can again be explained by meiotic recombination between monocentric circular chromatids leading to their catenation and formation of dicentric circles. It should be noted that results from earlier studies^17^ as well as our own WGS and PFGE analyses indicate that YJM311 has no systemic defects in meiotic recombination, as all chromosomes engage in seemingly normal meiotic recombination and crossover resolution (Fig. 5D and data not shown). Taken together, these results demonstrate that the circular structure of Chr1*b* is alone sufficient to explain the compromised meiotic viability seen in YJM311.

## Discussion

In this study, we used long read WGS and *de novo* genome assembly approaches to comprehensively define the structural genomic features of the heterozygous clinical yeast strain YJM311. Our construction of a haplotype-resolved diploid genome assembly exposed the complete architecture of YJM311 and enabled us to discern several unexplained characteristics of this strain, such as the clonal mosaicism within the stock population and the underlying cause of YJM311 spore inviability^14^. Even though this study analyzes the genomic structure of just a single diploid isolate of *S. cerevisiae*, our findings raise several important points which collectively argue for the integration of structural genomic information into models describing the origins, life histories, and evolution of strains and populations of *S. cerevisiae*.

Our conclusions indicate that structural genomic information will be very useful in identifying related individuals and lineages within a species. For example, because the *Z. bailii* circle (Fig. 3B) does not rely on surrounding homologous sequences for recombination-based integration^23^, it is very unlikely that identical insertions (*e*.*g*., into the right arm of Chr6) have occurred independently in distinct lineages of *S. cerevisiae*. Thus, this SV, which is easy to detect using long read WGS approaches, may be useful as an indicator of relatedness. Our own comparison of the *Z. bailii* circle integration site on Chr6 in YJM311 and another wild diploid isolate, JAY270, demonstrated that the site and orientation of insertion are identical between strains (data not shown). Thus, we suspect that the haploid parents harboring the *Z. bailli* circle-containing homologs of Chr6 in each strain may derive from a related lineage. Structural analysis of the *CUP1* array in wild isolates of *S. cerevisiae* is also likely to reveal new details about population relatedness and diversification in *S. cerevisiae*. Our finding that the Chr8*a CUP1* locus consists tandem Type 3 repeats suggests that the haploid parent of YJM311 which harbored Chr8*a* may be related to other strains harboring Type 3 arrays (*e*.*g*., YJM789)^21^. Moreover, *CUP1* array characterization not only informs on the evolutionary history of haploid strains, but it also sheds light onto the intra-genomic structural evolution that has occurred in the genomes of diploid strains. Future long read WGS surveys of SVs such as *Z. bailii* circle integrations and the *CUP1* locus in natural isolates of *S. cerevisiae* will undoubtedly provide an important and complimentary perspective of the population structure comprising this species.

Our study also highlights the importance of considering structural genomic information when drawing conclusions about the life history of a population or species. Previous studies have posited that diploid wild isolates of *S. cerevisiae* can be categorized into two classifications: 1) strains that sporulate efficiently and as a result undergo sexual cycles frequently such that they become homozygous, and 2) strains that sporulate inefficiently, rarely undergo sexual cycles, and thus, retain high levels of heterozygosity^8^. YJM311 does not fit neatly into either of these categories, as it sporulates proficiently yet remains highly heterozygous. One possible explanation for the retained heterozygosity of YJM311 is that although sporulates, the viability of resulting meiotic spores is greatly compromised by the circular nature of Chr1*b*. Thus, the overall efficacy of sexual reproduction in this strain is greatly diminished. Additionally, the poor spore viability of YJM311 is likely to lower the potential of intra-ascus mating events following spore germination, a mechanism of diploidization known to result in rapid homozygosis of the genome^29,30^.

Finally, our finding that the clonal mosaicism of YJM311 is caused by sub-telomeric alterations emphasizes the contributions of structural genomic variations to the diversification of a population. These regions of the genome have been largely invisible by short read WGS approaches, as the repetitive nature of sub-telomeric genes, Y’ elements, and telomere consensus sequences, precludes accurate assignment of reads to specific chromosomes. Yet, using long read WGS analysis, we and others have been able to interrogate these structures with increased rigor^31^. Together, our results demonstrate that these genomic regions are dynamic and that this plasticity reflects the activities of integral genomic stability pathways involved in homologous recombination and telomere maintenance^31-33^. As such, continued characterization of the structural attributes of telomeres in naturally evolved isolates will contribute a deeper understanding of the biology of telomeric structure and stability in *S. cerevisiae*. In this study, we have presented but a survey of the structural variations present within the YJM311 population, and have not yet conducted investigations to determine how these variations may differentially impact the fitness of YJM311 subclones in a natural or clinical setting. Future studies will build upon this work to determine if and how these SVs impart phenotypic change to cells, as well as to investigate how they promote continued genome evolution over time.

## Figure Legends

**Figure S1.**
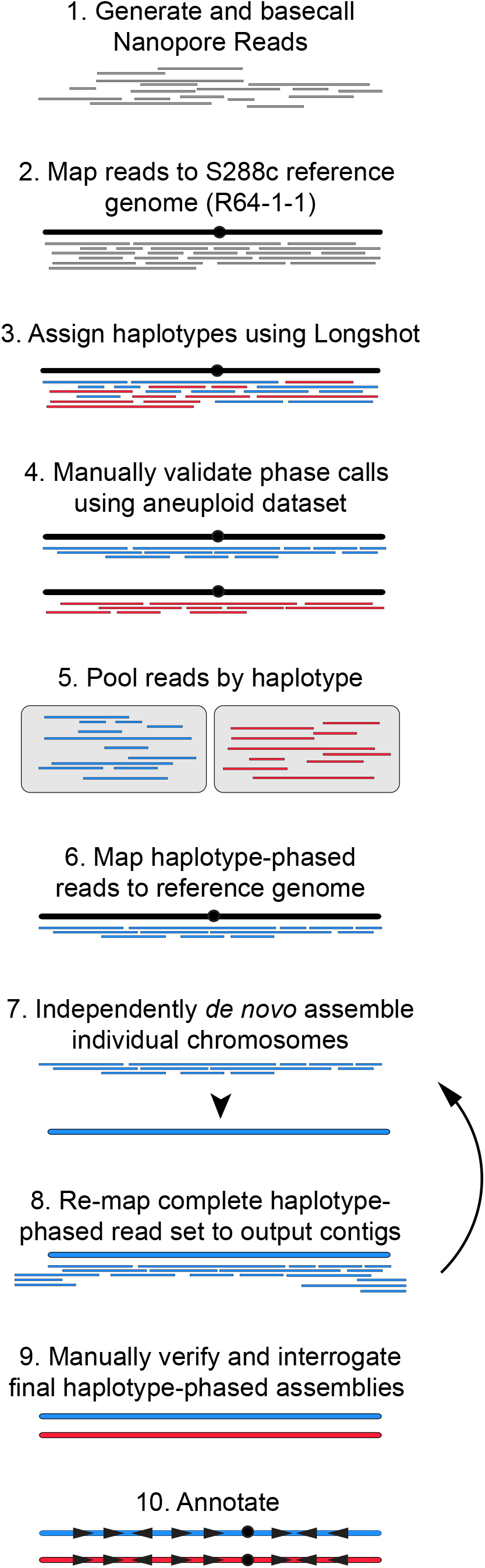
The *de novo* assembly pipeline used to construct the haplotype-phased diploid genome of YJM311-3. See Materials and Methods for detailed description.

**Figure S2.**
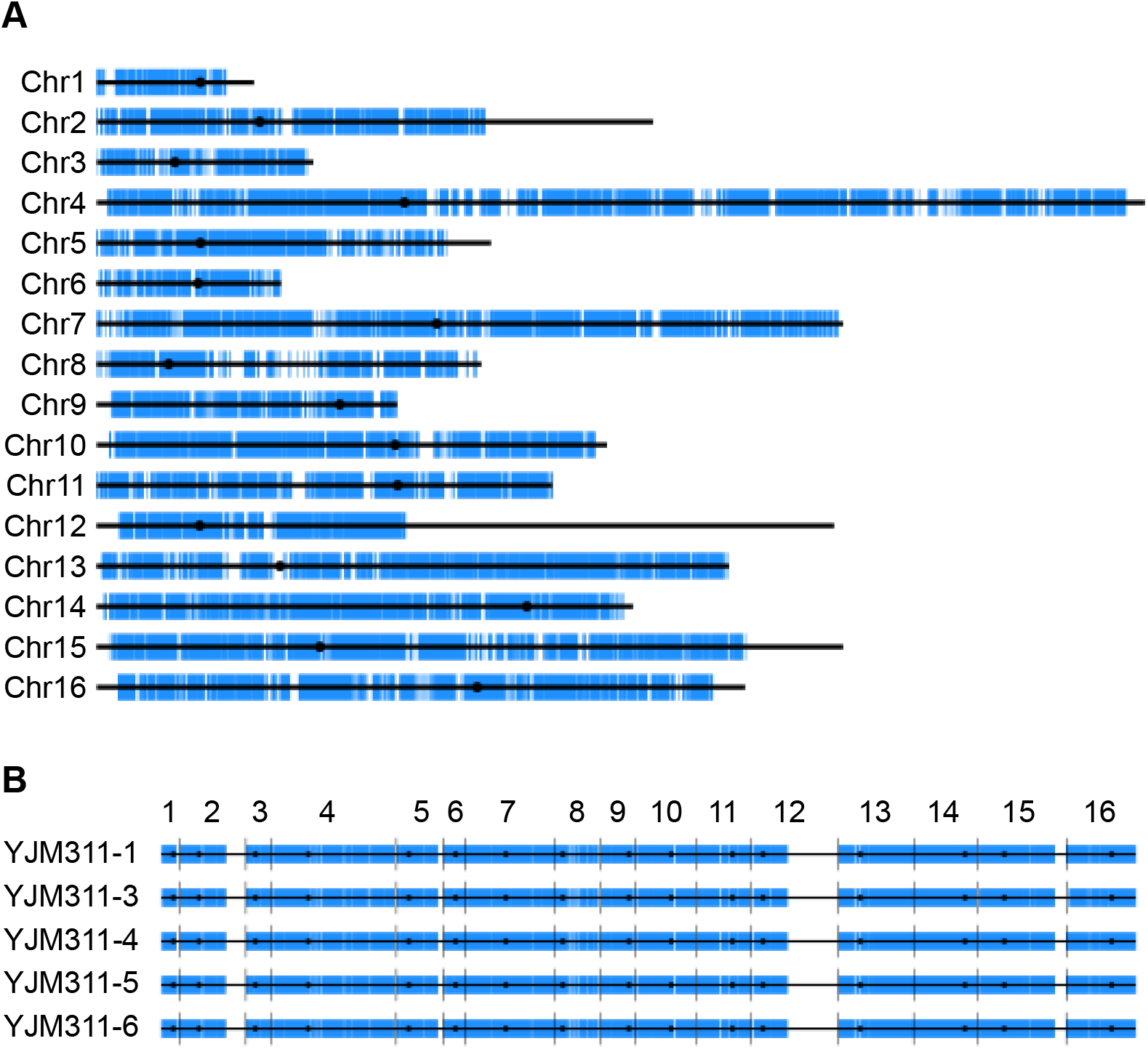
YJM311-derived subclones share identical hetSNP profiles. **A**. The high resolution hetSNP profile of the YJM311 genome organized by individual chromosome. **B**. The genomic hetSNP profiles of YJM311-1, YJM311-3, YJM311-4, and YJM311-5 shown together for comparative purposes.

## Materials and Methods

### Strains and Media

Our stock of YJM311 was a gift from Dr. Paul Magwene at Duke University. YJM311 subclones were isolated from this parental stock by streaking cells on a rich media YPD plate (10g/L yeast extract, 20 g/L Peptone, 20 g/L bacteriological agar, and 20g/L glucose) such that individual colonies could be recovered, and 7 independent colonies (YJM311-1 through YJM311-7) were picked at random for further characterization. For the sporulation analyses described in Fig. 5, YJM311-3 was streaked from a YPD plate to a sporulation media plate (10g/L potassium acetate, 1g/L yeast extract, 0.5g/L glucose, 0.35g/L complete drop out media, 20g/L bacteriological agar) and incubated at 30°C for 48 hours. Tetrads were digested with 25g/mL zymolyase on a YPD plate and dissected using a Zeiss dissecting microscope. Tetrad dissection plates were incubated for at 72 hours at 30°C prior to tetrad analysis.

### Pulsed Field Gel Electrophoresis (PFGE)

PFGE protocols and analysis were conducted as described previously^34,35^.

### Oxford Nanopore Minion Sequencing

To prepare reads for Oxford Nanopore Minion sequencing, high molecular weight DNA was isolated from cells, barcoded using the ONT ligation sequencing and native barcoding kits (SQK-LSK-109, EXP-NBD-104), and sequenced on a Minion flow cell (FLO-MIN106D). Reads were base called using ONT Guppy software and analyzed using CLC Genomics Workbench.

### Illumina sequencing

The genomes of the denoted YJM311 subclones and the 27 spores derived from YJM311-3 were sequenced using Illumina short read whole genome sequencing and analyzed as described previously^36^. Briefly, Genomic DNA from each clone was isolated using the Yeastar Genomic DNA kit from Zymo Research. A library of pooled, barcoded genomes was generated using a Seqwell plexWell-96 kit. The final barcoded library was sequenced using an Illumina HiSeq sequencer.

### De novo genome assembly

We began the assembly by first mapping long reads to the S288c reference genome (version R64-2-1) using CLC Genomics Workbench (Qiagen)(Fig. S1). We then phased the mapped reads by haplotype using the long read variant caller Longshot^19^. Longshot haplotype calling was performed for Minion reads as described at https://github.com/heasleyl/nanoporereads_haplotyping/blob/main/Pipeline. This analysis produced three distinct read groups designated as haplotype 1, haplotype 2, and reads whose haplotype could not be deduced. This last group of reads mapped primarily to the homozygous regions of the YJM311 genome (Fig. S2, A), so we combined this unphased group with each of the haplotype-phased read group to create two complete read sets with which to assemble chromosome-length scaffolds of each homologous chromosome.

We next verified that the Longshot haplotype calling was accurate. Our investigation of phenotype switching in YJM311 revealed that this behavior is primarily driven by stochastic and systemic genomic instability events resulting in the gain and loss of unique combinations of chromosomes^18^. As such, we had assembled a collection of YJM311-3 derived phenotypic variants, each of which harbored at least one aneuploidy. Aneuploidies alter the allele frequencies of each heterozygous single nucleotide polymorphism (hetSNP) site on the affected chromosome. For example, reduction in chromosome copy number from two copies to one copy results in chromosomal loss-of-heterozygosity (LOH) and the representation of only a single nucleotide identity at sites which had before been heterozygous. Thus, WGS analysis of these aneuploid YJM311-3 clones allowed us to deduce the hetSNP phasing of each chromosome and curate a list of genomic hetSNPs with the nucleotide identity resolved by haplotype. We then interrogated the nucleotide identities of these same hetSNPs in each Longshot haplotype-resolved long read set. We detected several phasing inaccuracies in the Longshot analysis, and these phase-switch errors coincided with genomic regions where YJM311-3 long reads broke alignment with the S288c reference genome due to heterozygous structural variations. We manually corrected these phasing errors, assembled new phase-confirmed haplotype-called long read sets, and proceeded with the genome assembly.

Manual browsing of the long reads indicated that the genome of YJM311-3 is highly syntenic with the reference genome. In general, structural divergence was limited to positions in the reference genome containing transposable element insertions, other repetitive loci such as the rDNA array and *CUP1* array, and sub-telomeric regions. With this in mind, we chose to assemble chromosome scaffolds individually using reads that partially or fully mapped to each reference chromosome using the ‘*de novo* assemble long reads’ function for CLC Genomics Workbench. This assembly method utilizes the open source programs minimap2 and miniasm^37,38^, and racon^39^. In many cases, this first round of assembly produced full length chromosomal contigs terminating with the consensus telomeric repeat sequences. However, any YJM311-3 reads which had not adequately mapped to the S288c reference genome would not have been incorporated into this first round of assembly. To ensure that all reads derived from YJM311-3 were utilized in the assembly of the genome, we performed an additional round of reassembly. For this second round of assembly, reads were remapped to the contigs produced from the first round and a new *de novo* assembly of each chromosome was performed. After this round, we confirmed the contiguity of each chromosomal assembly by manually inspecting the mapped reads for mis-mappings and breaks in alignment. At the end of this iterative assembly process, all reads from each haplotype-specific read set mapped accurately to each set of 16 *de novo* assembled contigs.

We annotated these contigs using the CLC-Genomics Genome Finishing Module ‘annotate from reference’ tool, which utilizes BLAST to identify annotations shared between a query sequence and an annotated reference (*i*.*e*., the S288c reference genome). We manually annotated Ty elements and sub-telomeric Y’ elements, as the program often failed to transfer annotations for these repetitive and degenerate sequences.

## Data and Materials Availability

Sequencing data and the annotated *de novo* assembly of YJM311 subclones and spores will be deposited on NCBI.

## Acknowledgements

We are grateful to Dr. Paul Magwene for sharing the strain YJM311. This study was supported by NIH/NIGMS awards 1K99GM13419301 to LRH and R35GM11978801 to JLA.

## Author Contributions

Conceptualization: LRH; Methodology: LRH; Investigation: LRH; Resources: LRH and JLA; Writing: LRH and JLA; Funding acquisition: JLA and LRH.

## Literature Cited

1 Peter, J. et al. Genome evolution across 1,011 Saccharomyces cerevisiae isolates. Nature, doi:papers3://publication/uuid/A69F80AD-26ED-4982-9809-8783E47F09F0 (2018).

2 Liti, G. et al. Population genomics of domestic and wild yeasts. Nature 458, 337–341, doi:10.1038/nature07743 (2009).

3 Peter, J. et al. Genome evolution across 1,011 Saccharomyces cerevisiae isolates. Nature 556, 339–344, doi:10.1038/s41586-018-0030-5 (2018).

4 Strope, P. K. et al. The 100-genomes strains, an S. cerevisiae resource that illuminates its natural phenotypic and genotypic variation and emergence as an opportunistic pathogen. Genome Res 25, 762–774, doi:10.1101/gr.185538.114 (2015).

5 Dutta, A. et al. Genome Dynamics of Hybrid Saccharomyces cerevisiae During Vegetative and Meiotic Divisions. G3 (Bethesda, Md.) 7, 3669–3679, doi:papers3://publication/doi/10.1534/g3.117.1135 (2017).

6 Fay, J. C. The molecular basis of phenotypic variation in yeast. Curr Opin Genet Dev 23, 672–677, doi:10.1016/j.gde.2013.10.005 (2013).

7 Bergström, A. et al. A high-definition view of functional genetic variation from natural yeast genomes. Mol Biol Evol 31, 872–888, doi:10.1093/molbev/msu037 (2014).

8 Magwene, P. M. et al. Outcrossing, mitotic recombination, and life-history trade-offs shape genome evolution in Saccharomyces cerevisiae. Proc Natl Acad Sci U S A 108, 1987–1992, doi:10.1073/pnas.1012544108 (2011).

9 Fischer, G., Liti, G. & Llorente, B. The Budding Yeast Life Cycle: more complex than anticipated? Yeast n/a, doi:https://doi.org/10.1002/yea.3533 (2020).

10 Peter, J. & Schacherer, J. Population genomics of yeasts: towards a comprehensive view across a broad evolutionary scale. Yeast 33, 73–81, doi:10.1002/yea.3142 (2016).

11 Almeida, P. et al. A population genomics insight into the Mediterranean origins of wine yeast domestication. Mol Ecol 24, 5412–5427, doi:10.1111/mec.13341 (2015).

12 McGinty, R. J. et al. Nanopore sequencing of complex genomic rearrangements in yeast reveals mechanisms of repeat-mediated double-strand break repair. Genome Res 27, 2072–2082, doi:10.1101/gr.228148.117 (2017).

13 Clemons, K. V., Hanson, L. C. & Stevens, D. A. Colony phenotype switching in clinical and non-clinical isolates of Saccharomyces cerevisiae. J Med Vet Mycol 34, 259–264, doi:10.1080/02681219680000441 (1996).

14 McCusker, J. H., Clemons, K. V., Stevens, D. A. & Davis, R. W. Genetic characterization of pathogenic Saccharomyces cerevisiae isolates. Genetics 136, 1261–1269 (1994).

15 McCusker, J. H., Clemons, K. V., Stevens, D. A. & Davis, R. W. Saccharomyces cerevisiae virulence phenotype as determined with CD-1 mice is associated with the ability to grow at 42 degrees C and form pseudohyphae. Infect Immun 62, 5447–5455, doi:10.1128/iai.62.12.5447-5455.1994 (1994).

16 Granek, J. A. & Magwene, P. M. Environmental and genetic determinants of colony morphology in yeast. PLoS Genet 6, e1000823, doi:10.1371/journal.pgen.1000823 (2010).

17 Granek, J. A., Murray, D., Kayrkçi, Ö. & Magwene, P. M. The genetic architecture of biofilm formation in a clinical isolate of Saccharomyces cerevisiae. Genetics 193, 587–600, doi:10.1534/genetics.112.142067 (2013).

18 Heasley, L. R. & Argueso, J. L. Systemic aneuploidization events drive phenotype switching in Saccharomyces cerevisiae. bioRxiv, 2021.2006.2016.448724, doi:10.1101/2021.06.16.448724 (2021).

19 Edge, P. & Bansal, V. Longshot enables accurate variant calling in diploid genomes from single-molecule long read sequencing. Nat Commun 10, 4660, doi:10.1038/s41467-019-12493-y (2019).

20 Sanchez, J. C. et al. Phenotypic and Genotypic Consequences of CRISPR/Cas9 Editing of the Replication Origins in the rDNA of. Genetics 213, 229–249, doi:10.1534/genetics.119.302351 (2019).

21 Zhao, Y. et al. Structures of naturally evolved CUP1 tandem arrays in yeast indicate that these arrays are generated by unequal nonhomologous recombination. G3 (Bethesda) 4, 2259–2269, doi:10.1534/g3.114.012922 (2014).

22 Borneman, A. R. et al. Whole-genome comparison reveals novel genetic elements that characterize the genome of industrial strains of Saccharomyces cerevisiae. PLoS Genet 7, e1001287, doi:10.1371/journal.pgen.1001287 (2011).

23 Galeote, V. et al. Amplification of a Zygosaccharomyces bailii DNA segment in wine yeast genomes by extrachromosomal circular DNA formation. PLoS One 6, e17872, doi:10.1371/journal.pone.0017872 (2011).

24 Sampaio, N. M. V. et al. Characterization of systemic genomic instability in budding yeast. Proc Natl Acad Sci U S A, doi:10.1073/pnas.2010303117 (2020).

25 Haber, J. E., Thorburn, P. C. & Rogers, D. Meiotic and mitotic behavior of dicentric chromosomes in Saccharomyces cerevisiae. Genetics 106, 185–205 (1984).

26 Murakami, H., Mu, X. & Keeney, S. How do small chromosomes know they are small? Maximizing meiotic break formation on the shortest yeast chromosomes. Curr Genet 67, 431–437, doi:10.1007/s00294-021-01160-9 (2021).

27 Murakami, H. et al. Multilayered mechanisms ensure that short chromosomes recombine in meiosis. Nature 582, 124–128, doi:10.1038/s41586-020-2248-2 (2020).

28 Mancera, E., Bourgon, R., Brozzi, A., Huber, W. & Steinmetz, L. M. High-resolution mapping of meiotic crossovers and non-crossovers in yeast. Nature 454, 479–485, doi:10.1038/nature07135 (2008).

29 Knop, M. Evolution of the hemiascomycete yeasts: on life styles and the importance of inbreeding. BioEssays: News and Reviews in Molecular, Cellular and Developmental Biology 28, 696–708, doi:papers3://publication/doi/10.1002/bies.20435 (2006).

30 Heasley, L. R., Singer, E., Cooperman, B. J. & McMurray, M. A. Saccharomyces spores are born prepolarized to outgrow away from spore-spore connections and penetrate the ascus wall. Yeast 38, 90–101, doi:10.1002/yea.3540 (2021).

31 Kockler, Z. W., Comeron, J. M. & Malkova, A. A unified alternative telomere-lengthening pathway in yeast survivor cells. Mol Cell 81, 1816-1829.e1815, doi:10.1016/j.molcel.2021.02.004 (2021).

32 Symington, L. S., Rothstein, R. & Lisby, M. Mechanisms and regulation of mitotic recombination in Saccharomyces cerevisiae. Genetics 198, 795–835, doi:papers3://publication/doi/10.1534/genetics.114.166140 (2014).

33 Wellinger, R. J. & Zakian, V. A. Everything you ever wanted to know about Saccharomyces cerevisiae telomeres: beginning to end. Genetics 191, 1073–1105, doi:10.1534/genetics.111.137851 (2012).

34 Zhang, H. et al. Gene copy-number variation in haploid and diploid strains of the yeast Saccharomyces cerevisiae. Genetics 193, 785–801, doi:10.1534/genetics.112.146522 (2013).

35 Argueso, J. L. et al. Double-strand breaks associated with repetitive DNA can reshape the genome. Proc Natl Acad Sci U S A 105, 11845–11850, doi:10.1073/pnas.0804529105 (2008).

36 Heasley, L. R., Watson, R. A. & Argueso, J. L. Punctuated Aneuploidization of the Budding Yeast Genome. Genetics 216, 43–50, doi:10.1534/genetics.120.303536 (2020).

37 Li, H. Minimap and miniasm: fast mapping and de novo assembly for noisy long sequences. Bioinformatics 32, 2103–2110, doi:10.1093/bioinformatics/btw152 (2016).

38 Li, H. Minimap2: pairwise alignment for nucleotide sequences. Bioinformatics 34, 3094–3100, doi:10.1093/bioinformatics/bty191 (2018).

39 Vaser, R., Sović, I., Nagarajan, N. & Šikić, M. Fast and accurate de novo genome assembly from long uncorrected reads. Genome Res 27, 737–746, doi:10.1101/gr.214270.116 (2017).

